# A Novel Registration Framework for Aligning Longitudinal Infant Brain Tensor Images

**DOI:** 10.1101/2024.07.12.603305

**Authors:** Kuaikuai Duan, Longchuan Li, Vince D. Calhoun, Sarah Shultz

## Abstract

Registering longitudinal infant brain images is challenging, as the infant brain undergoes rapid changes in size, shape and tissue contrast in the first months and years of life. Diffusion tensor images (DTI) have relatively consistent tissue properties over the course of infancy compared to commonly used T1 or T2- weighted images, presenting great potential for infant brain registration. Moreover, groupwise registration has been widely used in infant neuroimaging studies to reduce bias introduced by predefined atlases that may not be well representative of samples under study. To date, however, no methods have been developed for groupwise registration of tensor-based images. Here, we propose a novel registration approach to groupwise align longitudinal infant DTI images to a sample-specific common space. Longitudinal infant DTI images are first clustered into more homogenous subgroups based on image similarity using Louvain clustering. DTI scans are then aligned within each subgroup using standard tensor-based registration. The resulting images from all subgroups are then further aligned onto a sample-specific common space. Results show that our approach significantly improved registration accuracy both globally and locally compared to standard tensor-based registration and standard fractional anisotropy-based registration. Additionally, clustering based on image similarity yielded significantly higher registration accuracy compared to no clustering, but comparable registration accuracy compared to clustering based on chronological age. By registering images groupwise to reduce registration bias and capitalizing on the consistency of features in tensor maps across early infancy, our groupwise registration framework facilitates more accurate alignment of longitudinal infant brain images.

## Introduction

Brain image registration – the alignment of individual brain images to a standard brain image (i.e., template)— is important for establishing spatial correspondence and facilitating group-level inferences (Maintz & Viergever, 1998; Oliveira & Tavares, 2014). A number of approaches have been proposed for registering brain images, such as FMRIB’s Linear Image Registration Tool (FLIRT) (Jenkinson & Smith, 2001), FMRIB’s nonlinear image registration tool (FNIRT) (J. Andersson, Smith, & Jenkinson, 2008), Symmetric Normalization (SyN) algorithm (Avants, Epstein, Grossman, & Gee, 2008), Diffeomorphic Anatomical Registration Through Exponentiated Lie Algebra (DARTEL) (Ashburner, 2007) and its predecessor, Unified Segmentation (Ashburner & Friston, 2005) which is implemented in the Statistical Parametric Mapping (SPM) (Friston et al., 1994). While these algorithms have been successfully and routinely applied to register adult brain images, registering longitudinal infant brain images presents unique challenges (Evans, Janke, Collins, & Baillet, 2012; Shi et al., 2011). Over the course of infancy, the brain undergoes dramatic changes in size, morphology, myelination, and function (Chugani, 1998; Gao, Alcauter, Smith, Gilmore, & Lin, 2015; Huang et al., 2015; Pfefferbaum et al., 1994; Shi et al., 2011), with significant changes occurring in the infant’s brain almost every week (Devi, Chandrasekharan, Sundararaman, & Alex, 2015). Of particular relevance to registration, the relative signal intensities of gray and white matter in anatomical T1- and T2-weighted MRI images (the imaging modalities that are most commonly used for infant brain image registration (Dong, Wang, Lin, Shen, & Wu, 2017; D. Holland et al., 2014; Wei et al., 2022)) reverse over the course of the first postnatal months (Hayakawa, Konishi, Kuriyama, Konishi, & Matsuda, 1991; B. A. Holland, Haas, Norman, Brant-Zawadzki, & Newton, 1986; W. Zhang et al., 2015) (see supplementary Fig. S1). Given these rapid and substantial changes in tissue contrast and brain shape, infant brain images vary tremendously over developmental time, making it challenging to accurately identify and align corresponding brain features at different developmental stages.

A related challenge is the difficulty associated with selecting a template that is representative of the developmental variability within a longitudinal infant sample (Turesky, Vanderauwera, & Gaab, 2021). Selection of a representative template is critical in longitudinal studies because templates with features that are not well matched to the sample can introduce unnecessary deformations that may bias results (Evans et al., 2012; Guimond, Meunier, & Thirion, 2000). Although several age-specific pediatric templates have been created (Chen et al., 2022; Sanchez, Richards, & Almli, 2012; Shi et al., 2011), biases may still be introduced if the age-specific template is not closely matched to or equally representative of the age range under investigation (Van Hecke et al., 2011; Yoon, Fonov, Perusse, Evans, & Brain Development Cooperative, 2009). For instance, after creating an age-specific template for a relatively narrow age range (39- to 42-weeks gestational age), Kazemi et al. demonstrated that even narrower age range templates (39-40 and 41-42 weeks) improved registration performance(Kazemi, Moghaddam, Grebe, Gondry-Jouet, & Wallois, 2007). Given the fast pace of brain development in early infancy (with significant changes occurring on the order of days and weeks (Devi et al., 2015)) and the fact that individual infants develop on different time scales (with brain maturation unfolding more rapidly in some infants than others), registering infant images towards a sample-specific common space that is optimally representative of and specific to the sample of interest may yield more accurate registration than registering infant images to a predefined age-specific template (which may not be equally representative of all ages under investigation)(Kazemi et al., 2007).

To address the challenges associated with aligning highly heterogeneous longitudinal infant images, we developed and tested a novel approach for groupwise registration of diffusion tensor images (DTI) to a sample-specific common space. Our approach leverages 2 key registration insights —each reviewed below—to create a registration framework optimized for use in longitudinal infant research.

### 1. Registration of diffusion tensor images

Compared to T1 or T2-weighted images (which are most commonly used for infant brain image registration), tensor images offer relatively stable tissue properties over development (Dubois et al., 2021; G. Li et al., 2019; Tymofiyeva et al., 2013; Yoshida, Oishi, Faria, & Mori, 2013) and (unlike scalar maps) contain information about the microstructural orientation of white matter tracts which can be used to further differentiate brain structures, highlighting the potential of tensor images for more accurate alignment of infant brain images (see supplementary Fig. S1). While it has been shown that standard tensor-based registration (implemented in DTI-TK (https://dti-tk.sourceforge.net/)) outperforms scalar (fractional anisotropy or FA)-based registration for DTI scans of aging populations (H. Zhang et al., 2007; H. Zhang, Yushkevich, Alexander, & Gee, 2006), its performance in rapidly developing infant populations remains unexamined. Additionally, DTI-TK has not yet been implemented in a groupwise registration framework, an important consideration for registration of longitudinal infant brain images (as described below).

### 2. Groupwise registration

Groupwise registration is commonly used in infant neuroimaging studies to enhance registration accuracy (Ahmad et al., 2019; Dong, Cao, Yap, & Shen, 2019; Z. Tang & Fan, 2014; Q. Wang, Chen, Yap, Wu, & Shen, 2010). Groupwise registration first aligns subgroups of relatively more homogeneous images to their common spaces and then registers the resulting images to a final common space (Fig. 1B). A common implementation of this approach is to first align individual images to the closest predefined (age-matched) intermediary template (Chen et al., 2022; G. Li et al., 2015; Sanchez et al., 2012; Shi et al., 2011), and then align these intermediary templates to a final common space (Dubois et al., 2021; Gao et al., 2015; Jia, Yap, Wu, Wang, & Shen, 2011; Mangin et al., 2016; S. Tang, Fan, Wu, Kim, & Shen, 2009). Deformations estimated during this hierarchical process are then combined to transform each individual image from the native space to the final common space (S. Tang et al., 2009). (Mangin et al., 2016; Shi et al., 2011; S. Tang et al., 2009; Wu et al., 2015). While the advantages of groupwise registration are well documented (Jia et al., 2011; Lebenberg et al., 2018; S. Tang et al., 2009), it has not been implemented for registration of tensor-based images.

**Figure 1.**
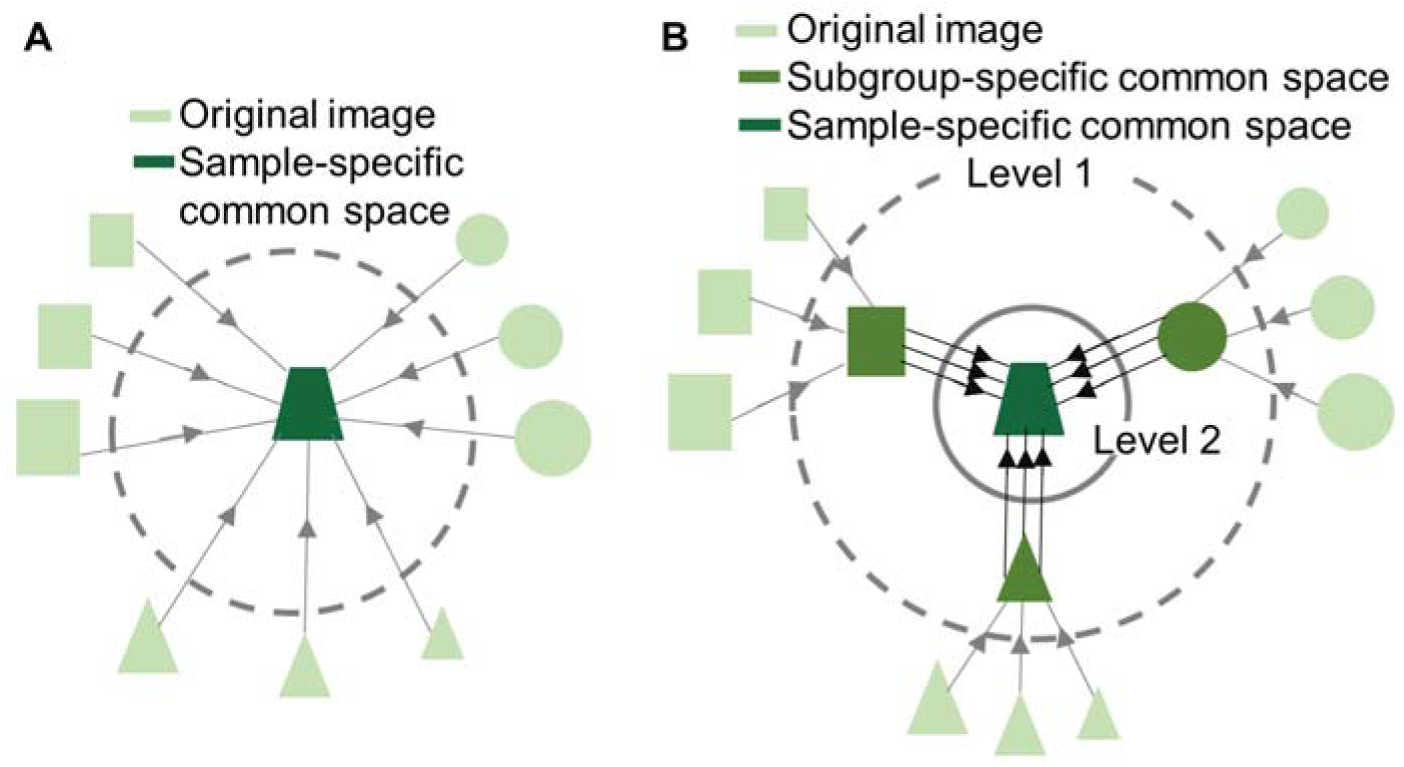
Illustration of (A) standard registration and (B) the proposed groupwise registration framework. In standard registration, all original images were directly aligned to a sample-specific common space. In the groupwise registration framework, original images were first clustered into subgroups based on shared image characteristics. In the 1st level, images within each subgroup were registered to their subgroup specific common space. In the 2^nd^ level, images that were aligned to the subgroup specific template in the 1^st^ level were further aligned to a sample-specific common space.

### 3. Our approach: Groupwise tensor-based registration to a sample-specific common space

To address the challenges associated with aligning highly heterogeneous longitudinal infant images, we leveraged the developmentally stable tissue properties and abundant white matter microstructural information within DTI images and the benefits of groupwise registration to a sample-specific common space. Unlike the standard tensor-based registration approach (DTI-TK), where all tensor images are registered together at one level to generate the sample-specific common space (Fig. 1A), our approach registers tensor maps at two levels, first registering a group of scans to a subgroup-specific space and then registering all resulting images to a sample-specific common space without assuming any predefined templates (Fig. 1B). We developed this registration approach using longitudinal infant DTI data collected from birth to 7 months, the most dynamic period of postnatal brain growth, providing a rigorous test case for evaluating our approach. Our aims were to: 1) replicate findings from aging populations demonstrating that standard tensor-based registration (DTI-TK) (H. Zhang et al., 2007) outperforms scalar (FA)-based registration in infants; and 2) compare the performance of our groupwise tensor-based registration approach with standard tensor-based registration (DTI-TK) (H. Zhang et al., 2007). Finally, we also compared the impact of 3 different approaches for clustering images into subgroups—clustering based on image similarity (which may yield more homogeneous subgroups, especially during periods characterized by rapid developmental change and/or individual differences in developmental timing), clustering based on chronological age (the predominant approach), and no clustering—on registration accuracy.

## 2. Materials and Methods

### 2.1. Participants

Participants were 27 typically developing infants (19 male and 8 female) enrolled in a prospective longitudinal study at the Marcus Autism Center, in Atlanta, GA, USA. Infants had a mean gestational age at birth of 39.09 weeks (SD = 1.40 weeks) and were considered to be typically developing on the basis of family history (no history of autism in up to 3^rd^ degree relatives and no history of developmental delay in 1^st^ degree relatives) and medical history (no pre- or perinatal complications, no history of seizures, no known medical conditions or genetic disorders, and no hearing loss or visual impairment). Each participant was scanned at up to three timepoints between birth and 7 months, for a total of 53 diffusion MRI scans. The distribution of participant age at each scan is displayed in Fig. S2. The Emory University Institutional Review board approved the research protocol for this study. Written informed consent was obtained from legal guardians for all infants included in this study.

### 2.2. Diffusion MRI data acquisition

All participants were scanned at the Center for Systems Imaging Core at Emory University School of Medicine using a 3T Siemens Trio Tim system with a 32-channel head coil. All infants were scanned during natural sleep, using the following procedure. First, infants were swaddled, rocked, and/or fed to encourage natural sleep. Once asleep, the infant was placed on a pediatric scanner bed. Scanner noise was reduced below 80 dBA by using: 1) sound attenuating pediatric headphones, equipped with MR-safe optical microphones to enable real-time monitoring of in-ear sound levels throughout the scan session; and 2) a custom-built acoustic hood, inserted into the MRI bore (Valente, Shultz, Klin, & Jones, 2014). To mask the onset of scanner noise, white noise—gradually increasing in volume—was played through the headphones prior to the first sequence. An MRI-compatible camera (MRC Systems) was mounted on the head coil to enable monitoring of the infant throughout the scan. A trained experimenter remained in the scanner room and the procedure was stopped if the infant awoke or if an increase in sound level was observed.

Structural, diffusion, and resting state MRI scans were collected at each session but only diffusion data were used in the current study. Diffusion MRI data were acquired using a multiband sequence (Feinberg et al., 2010; Moeller et al., 2010) with the following parameters: TR/TE of 6200/74ms, a multiband factor of 2 combined with a GRAPPA factor of 2, FOV of 184×184, Matrix of 92×92, b=0/700 s/mm^2^, spatial resolution of 2mm isotropic, 61 diffusion directions, 67 slices covering the whole brain, extra 6 averages of b0s. The total scan time for the diffusion MRI sequence was 7 minutes 26 seconds. A b0 image with the opposite phase encoding direction (posterior-to-anterior) was also collected for correcting the susceptibility-related distortion in diffusion MRI images (Glasser et al., 2013; L. Li, Rilling, Preuss, Glasser, & Hu, 2012).

### 2.3. Diffusion MRI data preprocessing

Diffusion MRI data were preprocessed using FSL 5.0.9 and in-house Matlab code (Matlab 2023). Preprocessing steps included correction for eddy-current distortion and removal of susceptibility distortion using the *eddy* and *topup* functions in FSL (J. L. Andersson, Skare, & Ashburner, 2003; J. L. R. Andersson & Sotiropoulos, 2016). Tensor maps and tensor-derived scalar maps, including maps of fractional anisotropy (FA) and mean diffusivity (MD), were calculated using FSL’s function *dtifit* with weighted least-square tensor fitting. Weighted least-square fitting was used to minimize the impact of motion on the infant data (Koay, Chang, Carew, Pierpaoli, & Basser, 2006).

### 2.4. Diffusion MRI data registration

Three different registration approaches—standard tensor-based registration, standard FA-based registration, and our novel groupwise tensor-based registration approach—were used and are described below.

#### 2.4.1. Standard tensor-based registration (DTI-TK)

Infants’ tensor maps were registered using the standard routine (https://dti-tk.sourceforge.net/pmwiki/pmwiki.php/Documentation.FirstRegistration) in DTI-TK (H. Zhang et al., 2007; H. Zhang et al., 2006). All participants’ tensor maps were first aligned to an initial target tensor template (generation of the initial target tensor template is described below) using a 6-degree of freedom (dof) rigid body transformation. Aligned images from all participants were then averaged to generate a 6-dof rigid body intermediate tensor template. This process was then repeated by aligning all participants’ tensor maps iteratively to the above-generated 6-dof rigid body intermediate tensor template via 12-dof affine transformations. These aligned images were then averaged to create a 12-dof affine intermediate tensor template. Lastly, all participants’ tensor maps were iteratively registered to the 12-dof affine intermediate tensor template using diffeomorphic transformation (via piecewise affine transformation that divides each image domain into uniform regions and transforms each region affinely) to generate the sample-specific common space.

##### Generation of the initial target tensor template

The initial target tensor template was generated by applying the abovementioned standard tensor-based registration method to align a subset of DTI scans (37 scans ranging from 0-7 months). A tensor map of an infant with relatively clear tissue contrast was chosen as the tensor template for 6-dof rigid body transformation (selection of tensor template did not affect the shape and size of the resulting initial target tensor template, see details in ***Effect of choosing different images for generating the initial target template*** in Supplementary Materials). This selected tensor template was nudged to closely match the origin of MNI space (the anterior commissure) and to be as straight as possible. Six-dof rigid body transformation, 12-dof affine transformation and diffeomorphic transformation were applied, as described above, to obtain diffeomorphically transformed tensor maps, which were then averaged to create the initial target tensor template for standard registration of tensor images. For standard registration of scalar FA images (see section 2.4.2), an FA map derived from the initial target tensor template was used as the initial target FA template.

#### 2.4.2. Standard registration of scalar FA images

FSL’s linear registration tool “FLIRT” and deformable registration tool “FNIRT” were used to align infant FA scalar images to a sample-specific common space. FNIRT is a medium-resolution nonlinear registration algorithm that has been previously used in developmental neuroimaging studies (Deniz Can, Richards, & Kuhl, 2013; O’Gorman et al., 2015; Westlye et al., 2010). The iterative registration approach for aligning all scans onto a sample-specific common space (Fig. 1A) is similar to the approach implemented in DTI-TK. Note that for each step of the registration, a single iteration was used as the majority of the literature do not employ multiple iterations for linear and non-linear registration for sample-specific scalar templates (Kazemi et al., 2007; L. Li et al., 2010; Sanchez et al., 2012).

#### 2.4.3. Groupwise tensor-based registration to a sample-specific common space

In our proposed groupwise registration framework, longitudinal infant DTI images are first clustered into more homogenous subgroups based on image similarity using Louvain clustering. DTI scans in each subgroup are then aligned separately using standard tensor-based registration (as in section 2.4.1). The resulting images from all subgroups are further aligned onto a sample-specific common space. These steps are described in detail below.

##### Defining similarity matrices among images

All participants’ tensor maps were first aligned to the initial target tensor template using 6-dof rigid body transformation. FA and medial diffusivity (MD) maps were then derived from the resulting tensor maps and used to compute pairwise distance between all aligned images (equations 1 and 2). These two DTI-derived metrics (FA and MD) were selected because FA maps differentiate between gray and white matter well, whereas MD maps show high contrast between brain tissue and cerebrospinal fluid (Jia et al., 2011). Next, the summed square of voxel-wise differences of FA and MD maps between each image pair were calculated and normalized to range between 0 and 1, and then used to compute the similarity index between image A and B as in equations 1-3.

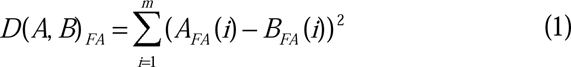

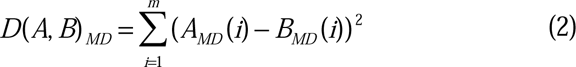

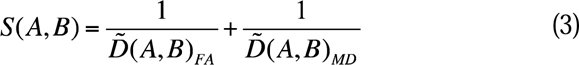

where *m* is the total number of voxels. *A_FA_*(*i*) and *B_FA_*(*i*) refer to the FA values of images *A* and *B* at brain voxel *i*. *D*(*A, B*)*_FA_* is the distance of *FA* maps between images *A* and *B*. *A_MD_* (*i*) and *B_MD_*(*i*) refers to the MD values of images *A* and *B* at brain voxel *i*. *D*(*A, B*)*_MD_* is the distance of MD maps between images *A* and *B*. *D̃*(*A, B*)*_FA_* and *D̃*(*A, B*)*_MD_* represent the normalized (values between 0 and 1) FA and MD distance value between images *A* and *B*. *S*(*A, B*) is the similarity index between images *A* and *B*.

##### Grouping scans based on image similarity using Louvain clustering

To stratify scans into more homogeneous subgroups based on their image similarity, we applied Louvain clustering to the similarity matrices estimated using FA and MD maps. Louvain clustering maximizes within-group connections and minimizes between-group connections (Reichardt & Bornholdt, 2006; Vincent D Blondel, Jean-Loup Guillaume, Lambiotte, & Lefebvre, 2008). The number of groups was determined in an iterative manner, starting with a negative initial modularity score and with one cluster. The partition process was repeated until the observed increase in modularity score was less than a given threshold (1E-9) (Blondel, Guillaume, Lambiotte, & Lefebvre, 2008).

##### Groupwise tensor-based registration

After the scans were clustered into more homogeneous subgroups based on their shared image features, level 1 registration was performed within each subgroup using standard tensor-based registration (described in section 2.4.1): the original tensor maps in each subgroup were aligned to their respective common space to generate subgroup specific tensor templates via rigid body transformation, affine and deformable transformations. In level 2 registration, the subgroup specific tensor templates from each subgroup were then aligned onto the sample-specific common space using standard tensor-based registration (Fig. 1B). Finally, the original tensor maps were transformed from each individual’s original space to the sample-specific common space via the transformations derived in the two-level alignment process.

##### Effect of clustering strategy on groupwise tensor-based registration

To compare the effect of different clustering strategies on registration performance, we also considered (1) subgrouping scans based on chronological age; and (2) no clustering (treating all scans as a single group and performing two rounds of standard tensor-based registration on all scans). Registration performance following each clustering strategy was compared against that from Louvain clustering based on image similarity.

##### Effect of brain masks and number of iterations in groupwise tensor-based registration

To evaluate whether the proposed groupwise tensor-based registration was robust to different brain masks, we compared registration performance when brain masks were selected with FA thresholds of 0.05 (i.e., whole-brain), 0.1 (white matter-enriched and some gray matter regions), and 0.25 (white matter heavy regions). Moreover, we examined the effect of varying the number of iterations in affine and deformable transformation stages.

#### 2.4.4. Registration performance evaluation

We employed three commonly used metrics to evaluate the performance of different registration methods. The first metric is dyadic coherence, κ, which quantifies the variability in the aligned principal eigenvectors across scans (Basser & Pajevic, 2000; Jones et al., 2002). Dyadic coherence ranges from 0 to 1, with 0 for randomly oriented tensor directions and 1 for perfectly aligned tensors in each image voxel across scans. The second metric is the voxel-wise normalized standard deviations across all FA (σ_FA_) maps, which was computed for each voxel within the FA mask. (H. Zhang et al., 2006). Suboptimal alignment strategies that overlap different white matter structures onto each other are expected to have high normalized standard deviation of FA. When plotting the empirical cumulative distribution functions (CDF), methods with better alignment are expected to have CDFs of κ and σ_FA_ to the right and left, respectively. The third metric is the normalized mutual information (NMI) value (Studholme, Hill, & Hawkes, 1999) between each FA map of the aligned tensor map and the average FA map across all aligned tensor maps, which is computed by dividing their joint entropy by the sum of the marginal entropies. NMI values range from 0 to 1 and reflect the similarity between each aligned FA map and their average. Larger NMI values indicate higher similarity (i.e., better registration) between each aligned scan and their average. Pairwise two sample t-tests were used to compare the NMI values from different registration methods. Unless noted, all statistics were computed for brain voxels with FA>0.25 in the average FA map that was rigidly aligned to the sample-specific common space for fair comparison. Maps of σ_FA_ were also generated and compared to evaluate the performance of the different registration methods.

## 3. Results

### Standard FA-based registration vs. standard tensor-based registration

Figs. 2A, 2C and 2D plot the performance of standard FA-based registration and standard tensor-based registration. Standard tensor-based registration generated smaller σ_FA_ than FA-based registration (Fig. 2A and the bottom row of Fig. 2D) for both affine and deformable transformation stages, confirming previous findings in aging populations (H. Zhang et al., 2007). Moreover, standard tensor-based registration achieved significantly larger NMI values than FA-based registration (Fig. 2C, *p* < 1e-16), indicating the similarity of FA maps derived from aligned tensor maps using standard tensor-based registration was significantly higher than that from standard FA-based registration. The mean and SD of FA maps from standard FA-based registration generated overall higher variability in gray matter than standard tensor-based registration, resulting in less well-defined gyri and sulci boundaries (Fig. 2D). Overall, standard tensor-based registration outperformed standard FA-based registration by generating less variable and more similar FA maps across infant longitudinal DTI scans.

**Figure 2.**
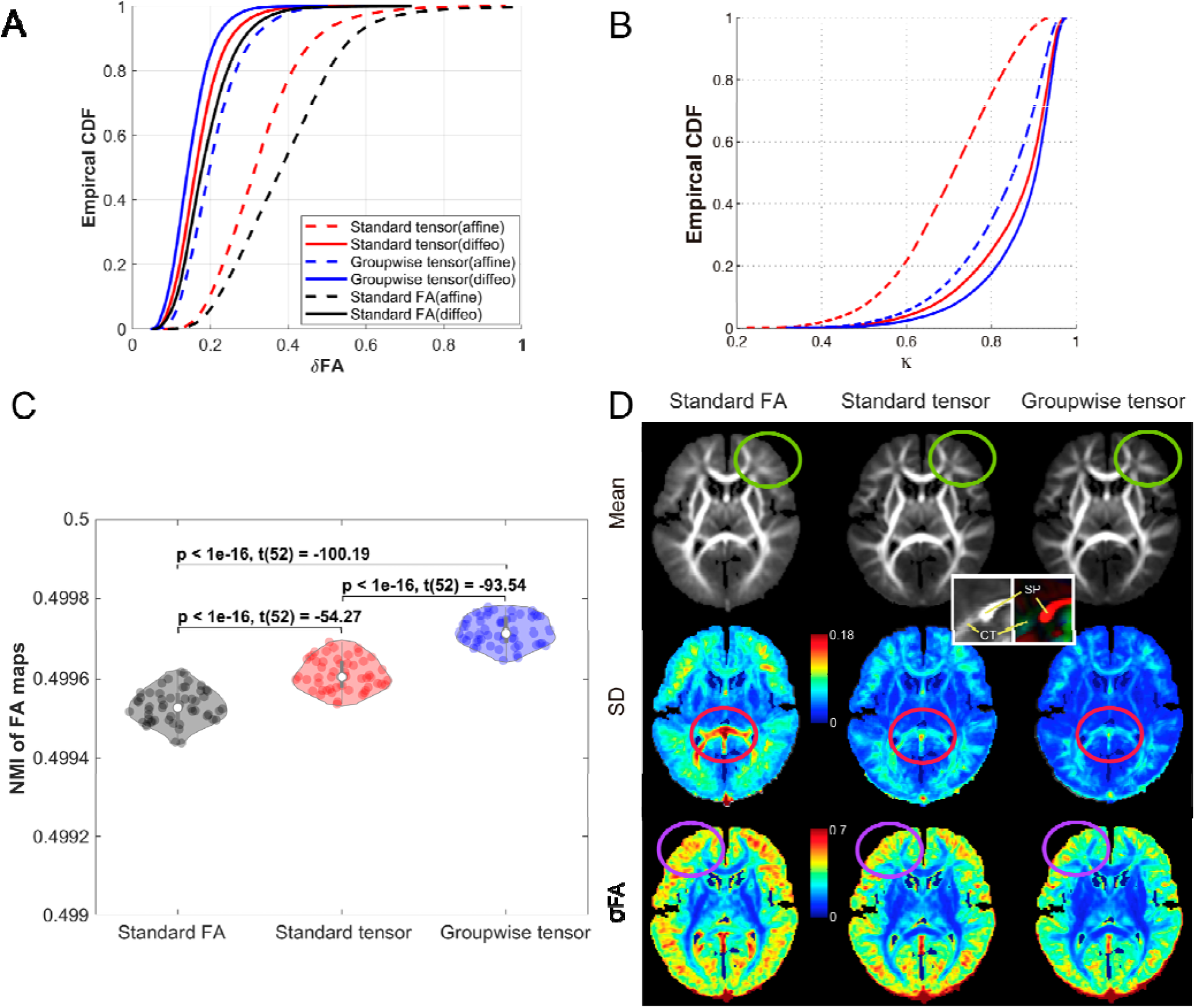
Registration performance of standard FA-based registration, standard tensor-based registration, and groupwise tensor-based registration. (A) CDF plots of normalized standard deviation of FA maps (σ_FA_) obtained from standard FA-based registration (black), standard tensor-based registration (red), and groupwise tensor-based registration (blue). For each of the registration method, dashed line represents the results from the affine (linear) transformation stage and solid line represents the results from the diffeomorphic (diffeo, nonlinear) transformation stage (the same for subplot (B)). (B) CDF plots of dyadic coherence (κ) derived from standard tensor-based registration (red) and groupwise tensor-based registration (blue). Dyadic coherence was not evaluated for FA-based registration since tensor information was not used during FA-based registration. (C) NMI values and pairwise two-sample t-test statistics of FA maps derived from FA-based registration (black), standard (red) and groupwise (blue) tensor-based registration. Groupwise tensor-based registration generated the highest registration accuracy (as reflected by the smallest σ_FA_, the largest dyadic coherence, and the largest NMI values), followed by standard tensor-based registration, and then standard FA-based registration. (D) Mean, SD and σ_FA_ of voxels within FA maps derived using each approach. Voxels within FA maps derived using standard FA-based registration show greater variability compared to (standard or groupwise) tensor-based registration methods, especially at the splenium of the corpus callosum (magenta circles), leading to less well-defined gyri and sulci boundaries (green and purple circles).

### Standard tensor-based registration vs. groupwise tensor-based registration

In groupwise tensor-based registration, three subgroups (Fig. S3A) were identified using Louvain clustering based on image similarity with the default resolution value (i.e., resolution = 1). As expected, cluster membership was correlated with scan age, brain volume, and mean FA, but not driven by any single metric (Fig. S3B and Fig. S4).

Compared to standard 1-level tensor-based registration, groupwise (2-level) tensor-based registration yielded smaller σ_FA_ (Fig. 2A and the bottom row of Fig. 2D), larger dyadic coherence (Fig. 2B), and significantly larger NMI values for FA maps (Fig. 2C, *p* < 1E-16, i.e., significantly higher similarity between the aligned FA maps and their group average derived from all the aligned DTI scans). These differences in registration accuracy were observed in both affine and deformable alignment stages, but were particularly pronounced in the affine stage (Figs. 2A and 2B), suggesting that the alignment to the subgroup specific space during the first level registration may be critical for the improved accuracy in the 2^nd^ level registration.

A closer look at the maps of the standard deviation and σ_FA_ generated using the three registration methods revealed that the splenium of the corpus callosum from standard FA-based registration yielded especially high σ_FA_ values when compared to those from (standard or groupwise) tensor-based registration methods (Fig. 2D, magenta circles). Examination of structures (Fig. 2D, inset) in this region illustrates that differentiation between the splenium and the cerebellar tentorium—both of which have high FA values—can be achieved when using the distinct orientation information available in tensor, but not FA, maps.

### Effect of different clustering strategies in groupwise tensor-based registration

#### (1) Clustering by image similarity vs. by chronological age

Longitudinal infant scans were clustered into 3 subgroups based on their chronological age: group 1: age <3 months (21 scans); group 2: age ≥ 3 and age < 6 months (25 scans); group 3: age ≥ 6 months (7 scans). For groupwise tensor-based registration, clustering by image similarity or by chronological age yielded comparable registration performance: CDF plots of dyadic coherence and σ_FA_ are largely overlapping (Fig. 3A and 3B) and MNI values of FA maps were not significantly different between clustering approaches (Fig. 3C, *p* = 0.24, *t* (52) = 1.19). This lack of difference may be explained by the relatively large amount of overlap between subgroups generated by each approach: 18 out of 26 scans (69%) in subgroup 1 clustered by image similarity were between 3 and 6 months of age; 17 out of 22 scans (77%) in subgroup 2 clustered by image similarity were between 0 and 3 months of age (Fig. 3D).

**Figure. 3.**
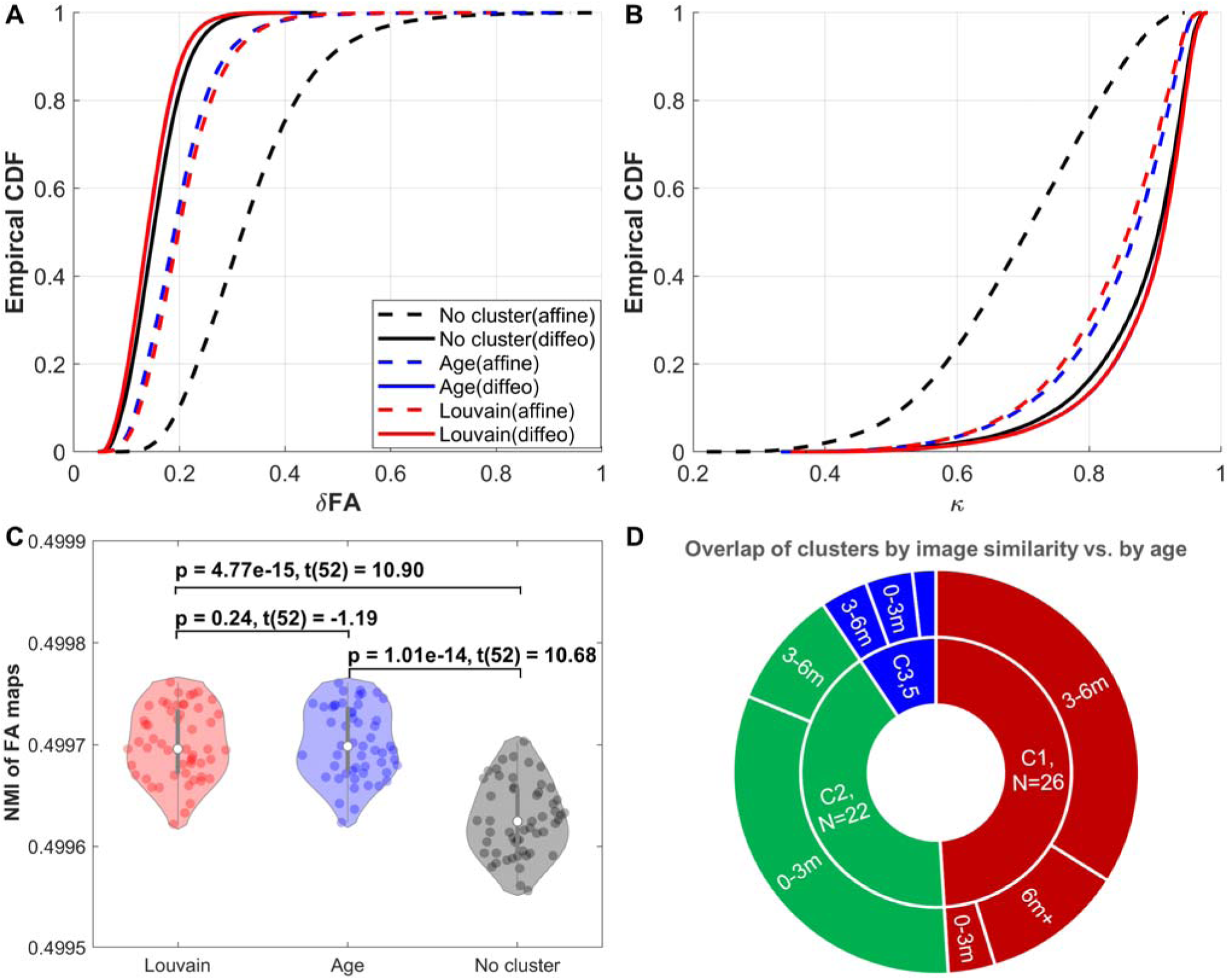
CDF plots of (A) normalized standard deviations of FA (σ_FA_) and (B) dyadic coherence (κ) derived from groupwise tensor-based registration clustered by image similarity (Louvain, red lines) or chronological age (Age, blue lines) or without clustering (No cluster, black lines). For each of the clustering methods, dashed lines represent the results from the affine (linear) transformation stage and solid lines represent the results from the diffeomorphic (diffeo, nonlinear) transformation stage (the same for subplot (B)). (C) NMI values and pairwise two-sample t-test statistics of FA maps derived from groupwise tensor-based registration clustered by image similarity (Louvain, red) or chronological age (Age, blue) or without clustering (No cluster, black), and (D) Overlap between clusters generated by image similarity (C1-C3, the inner circle) and by chronological age (the outer circle, where 0-3m denotes age <3 months, 3-6m represents age ≥ 3 and age < 6 months, and 6m+ denotes age ≥ 6 months). Compared to no clustering, clustering by image similarity or by chronological age yielded significantly improved registration accuracy, as indexed by smaller σ_FA_ and significantly larger NMI values. This indicates that creation of relatively homogenous subgroups on the basis of shared features (image similarity or age) is critical for improving accuracy in groupwise tensor-based registration. For groupwise tensor-based registration, clustering by image similarity or by chronological age yielded similar registration performance as indicated by largely overlapping CDF plots, possibly due to the large overlap in clusters resulting from the two clustering methods.

#### (2) Clustering based on image similarity vs. no clustering

Performing groupwise tensor-based registration without clustering (treating all 53 longitudinal infant DTI scans as a single group) did not provide comparable registration accuracy at both affine and diffeomorphic transformation stages compared to clustering by image similarity (Fig. 3(A)-(C)), indicating that the improved accuracy of groupwise tensor-based registration compared to standard tensor-based registration is not simply driven by running two rounds of affine and deformable registrations (i.e., initial clustering of images into more homogeneous subgroups is critical).

### Effect of varying brain masks and varying the number of iterations in groupwise tensor-based registration

Groupwise tensor-based registration outperformed standard tensor-based registration across a range of masking approaches (see details in ***Effects of brain masks with different FA thresholds*** in Supplementary Materials). Moreover, increasing the number of iterations to register individual tensor images during the affine and diffeomorphic transformation stages increased registration accuracy, but the improvement was negligible when the number of iterations was greater than 2 (see details in ***Effects of increasing number of iterations in groupwise tensor-based registration*** in Supplementary Materials).

## 4. Discussion

Building on previous research demonstrating the advantages of tensor-based over scalar-based registration (Y. Wang et al., 2016; H. Zhang et al., 2007; H. Zhang et al., 2006; Hui Zhang, Yushkevich, Rueckert, & Gee, 2009) and the benefits of groupwise over standard registration (Jia et al., 2011; Lebenberg et al., 2018; S. Tang et al., 2009), we developed a groupwise tensor-based registration framework for aligning longitudinal infant brain images collected between birth and 7 months, a period marked by very rapid postnatal brain growth and change. Briefly, longitudinal infant DTI maps were first clustered into several smaller and homogenous subgroups based on image similarity using Louvain clustering. Then, standard tensor-based registration was implemented groupwise: first to align all tensor images within a subgroup to their subgroup-specific common space, and then to register the images in the subgroup common space to the sample-specific common space.

Compared to scalar (FA)-based registration, both standard and groupwise tensor-based registration improved registration accuracy globally (as quantified by smaller normalized standard deviations of FA, and larger NMI values between aligned FA images and their average) and locally (as indicated by more sharply defined gyri and sulci boundaries, especially in the splenium of the corpus callosum), confirming that differentiation of distinct brain structures with similar anisotropic FA values can be achieved using the orientation information embedded in tensor maps, but not FA maps (Y. Wang et al., 2016; H. Zhang et al., 2007).

Compared to both standard FA-based and standard tensor-based registration, groupwise tensor-based registration significantly improved registration accuracy globally, as quantified by larger dyadic coherence of the principal eigenvector of the tensor maps, smaller normalized standard deviations of FA, and larger NMI values between aligned FA images and their average. These improvements in registration accuracy were observed in both affine and deformable alignment stages, but were particularly pronounced in the affine stage, suggesting that 1^st^-level registration of tensor images within subgroups may be critical for the improved accuracy in the affine stage at the 2^nd^ level registration.

Importantly, clustering of images into subgroups impacted groupwise registration accuracy. Clustering based on image similarity and clustering based on chronological age outperformed no clustering (i.e., applying standard tensor-based registration to all images twice), suggesting that creating more homogeneous subgroups with shared features (in this case, image similarity or chronological age) is critical for yielding improved registration performance. Contrary to our initial predictions, clustering based on image similarity did not improve registration accuracy compared to clustering based on chronological age, a null result that may be explained by the largely overlapping subgroups generated by each approach. Given that clustering by image similarity performs as well as clustering by chronological age, it may be advantageous to adopt the former approach when faced with uncertainty about which age cutoffs are most likely to yield homogeneous subgroups, or when working across developmental periods characterized by pronounced individual differences in developmental timing (with some infants maturing on different time scales than others).

While the present study focused primarily on registration of diffusion scans, our approach can also be useful for registering infant anatomical and functional images collected within the same session as diffusion scans. Specifically, the warping/deformation fields derived from groupwise registration of DTI maps can be readily applied to anatomical and functional images collected in the same session, potentially providing more accurate alignment for these imaging modalities especially during early infancy when gray and white matter tissue contrasts are isointense in T1- and T2-weighted images. Additionally, while we developed this registration approach using data collected from birth to 7 months—a particularly dynamic period of growth, providing a rigorous test case for this approach—this registration strategy can be readily applied to infants outside of our age range and can accommodate a variety of longitudinal sampling designs.

It should be noted while sample-specific templates have the advantage of minimizing deformations between individual images and the sample-specific common space, sample-specific templates usually lack standardized stereotaxic coordinates, making stereotaxic mapping and cross-study comparisons challenging (Evans et al., 2012). A potential solution is to report findings based on brain regions (instead of coordinates) using standard parcellations (Akiyama et al., 2013; Shi et al., 2011). If coordinate-based reporting of results is necessary, a transformation from the sample-specific template to standard stereotaxic space can be performed (Andersen et al., 2005; Chen et al., 2022; Shi et al., 2011).

As research increasingly focuses on mapping trajectories of brain development during infancy (Deoni et al., 2022; Edwards et al., 2022; Fitzgibbon et al., 2020; Howell et al., 2019)—a highly dynamic period that likely exerts a strong influence on neurodevelopmental disorders (Shen & Piven, 2017)—there is a growing need for methodological tools designed to address the unique challenges inherent to longitudinal infant neuroimaging research. Here we present a novel groupwise tensor-based registration approach (made publicly available at https://github.com/Luckykathy6/groupwiseRegister), specifically designed to address challenges inherent to registration of rapidly changing longitudinal infant brain images. Given that accurate alignment of brain structures across participants is a cornerstone for atlas-based analyses of developing brains, we believe that the proposed method will aid in advancing understanding of early brain development, a critical imperative for supporting children with neurodevelopmental disorders.

## Acknowledgement

We greatly appreciate the families and their infants who volunteered to participate in this research study. We would also like to thank the research coordinators at the Marcus Autism Center, Brittney Sholar, Carly Reineri, and Joanna Beugnon, and Michael White, the MRI Tech at the Emory Center for Systems Imaging Core, for leading data collection efforts, as well as Dr. Lei Zhou and Michael Valente for their assistance with equipment development and data acquisition protocols, Mahmoud Zeydabadinezhad for his help with data processing, and Zening Fu for his critical comments on the manuscript. The work is funded by the National Institutes of Mental Health, USA (K01MH108741, 2P50MH100029, and R01MH119251 to SS); the National Institute of Biomedical Imaging and Bioengineering, USA (R01EB027147 to SS and VC); and funds from the Whitehead and Marcus Foundations (to SS).

## Data and Code Availability

All scripts are available in the public repositoryhttps://github.com/Luckykathy6/groupwiseRegister. The DTI data will be made available on Zenondo or OpenNeuro once the paper is published.

## Author contributions

K. D., L. L. and S. S. conceptualized the study. K. D. and L. L. performed data analysis. K. D., L. L. and S. S. wrote the manuscript. S. S., L. L. and V. D. helped interpret the results and revise the manuscript. S. S., L. L. and V. D. contributed to funding acquisition.

