## Supplementary material for "A Novel Registration Framework for Aligning Longitudinal Infant Brain Tensor Images"

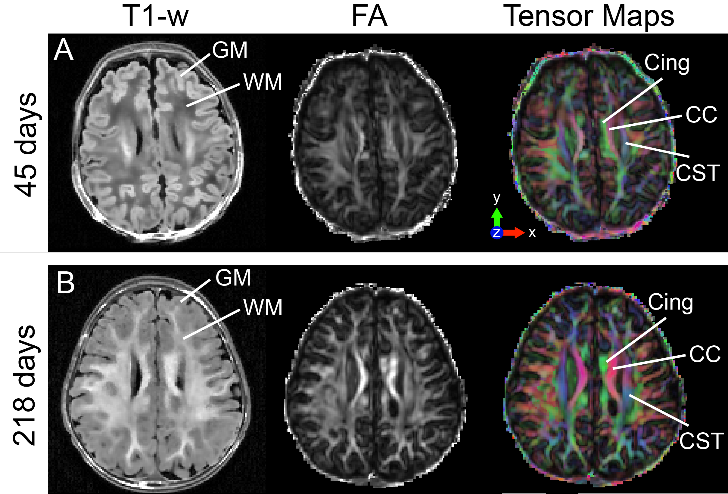

**Figure S1**. T1-weighted (T1w), fractional anisotropy (FA), and map of principal eigenvector of diffusion tensor images from (A) a 45-day-old infant and (B) a 218-day-old infant. T1-weighted (T1-w) images illustrate the developmental variability in infant brain images, where gray matter is brighter than white matter in (A) younger infants, and reversed in (B) older infants. By contrast, FA and tensor maps show similar patterns of tissue contrast across age. Tensor maps provide orientation information about WM microstructure, allowing for more detailed mapping of features between individuals as illustrated by their capability to identify structures such as the cingulum (cing), corpus callosum (CC), and corticospinal tract (CST) using the distinct orientations of these pathways.

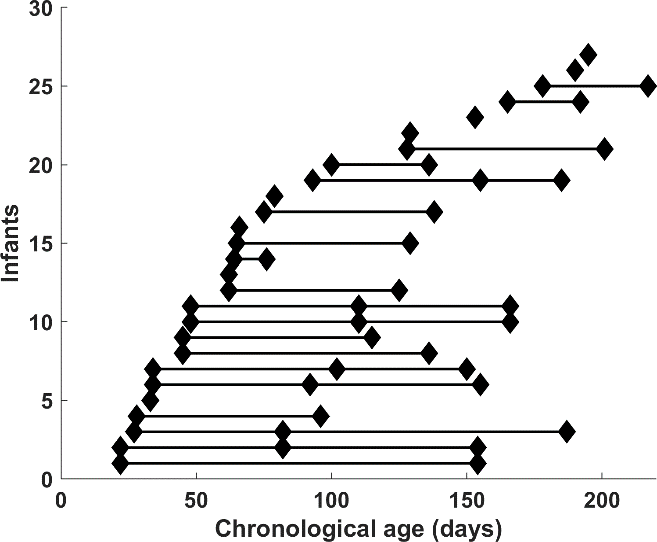

**Figure S2**. Included infants and their scan ages. Infants were scanned up to 3 scans between birth and 7 months, using a randomized, non-uniform longitudinal sampling design.

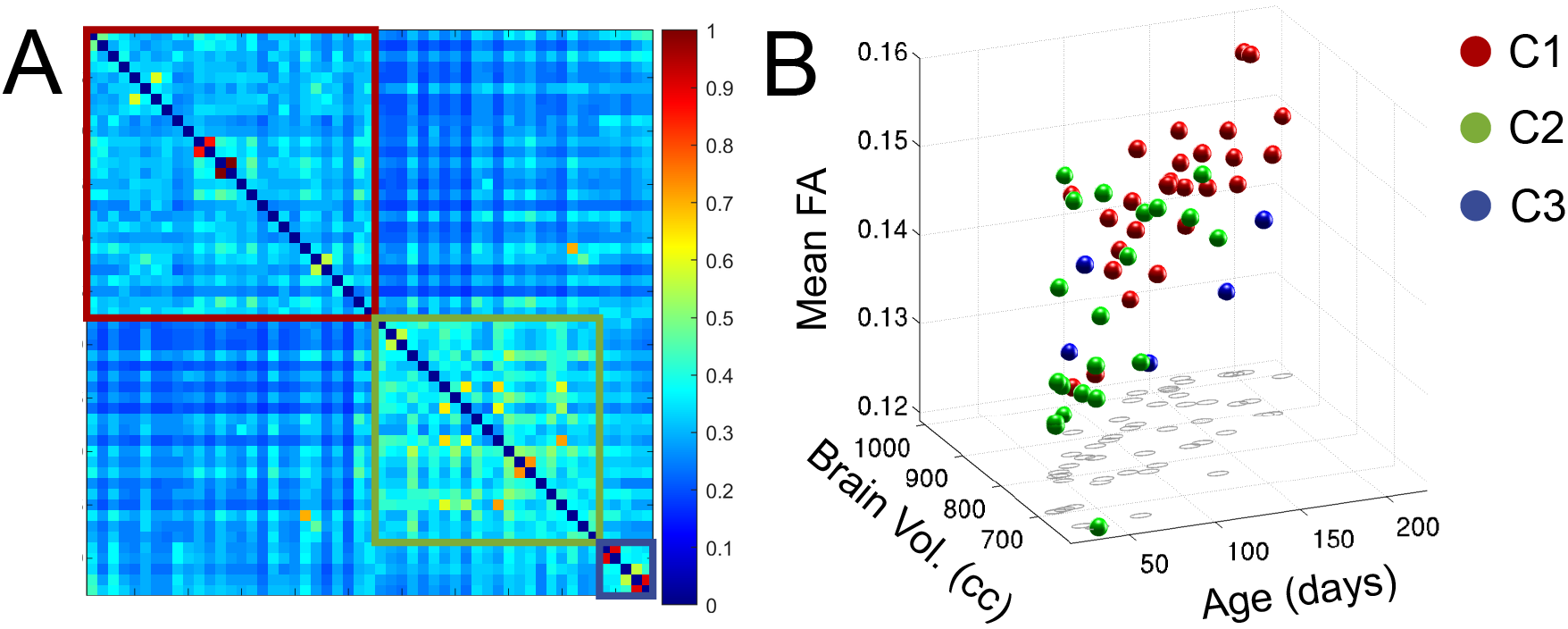

**Figure S3**. Clustering results and subgroup characteristics. (A) The similarity matrix for the 53 infant longitudinal DTI scans. Higher values (warmer colors) indicate higher similarity between two scans. The 3 subgroups detected by Louvain clustering are outlined by boxes with different colors to indicate their memberships (cluster 1: N = 26 scans, cluster 2: N = 22 scans, cluster 3: N = 5 scans). (B) The relationship between subgroup membership, mean FA, brain volume and age.

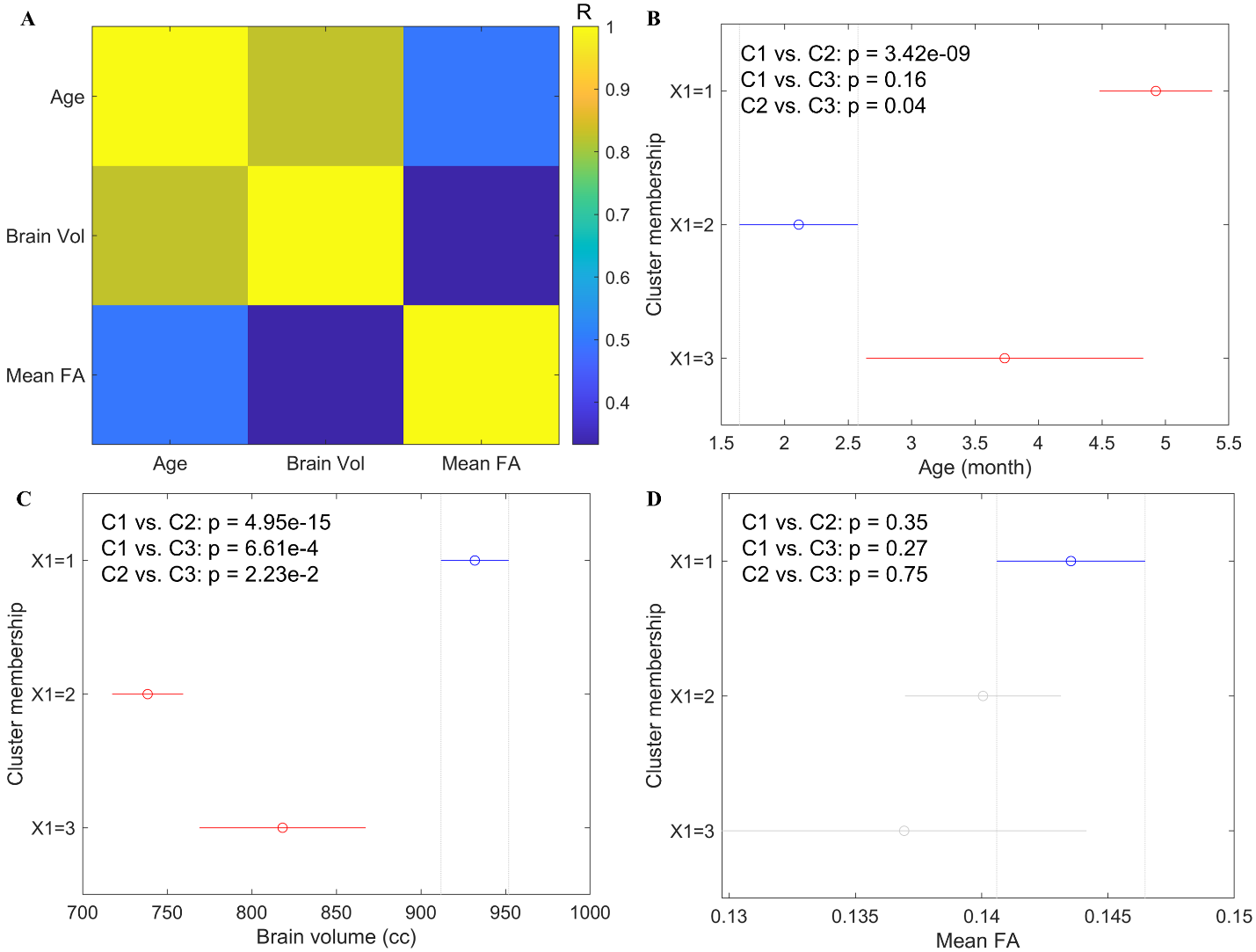

**Figure. S4**. (A) Cross-correlation among age, brain volume and mean FA. Pairwise difference of (B) age, (C) brain volume and (D) mean FA among the 3 subgroups.

***Effects of brain masks with different FA thresholds.*** Groupwise tensor-based registration showed substantially improved registration accuracy (larger κ and smaller σ_FA_) compared to standard tensor-based registration (Fig. S5 (A)) at three different brain mask levels (Fig. S5 (B), FA > 0.05: whole-brain; FA > 0.1: white matter-enriched and some gray matter regions; FA > 0.25: white matter heavy regions).

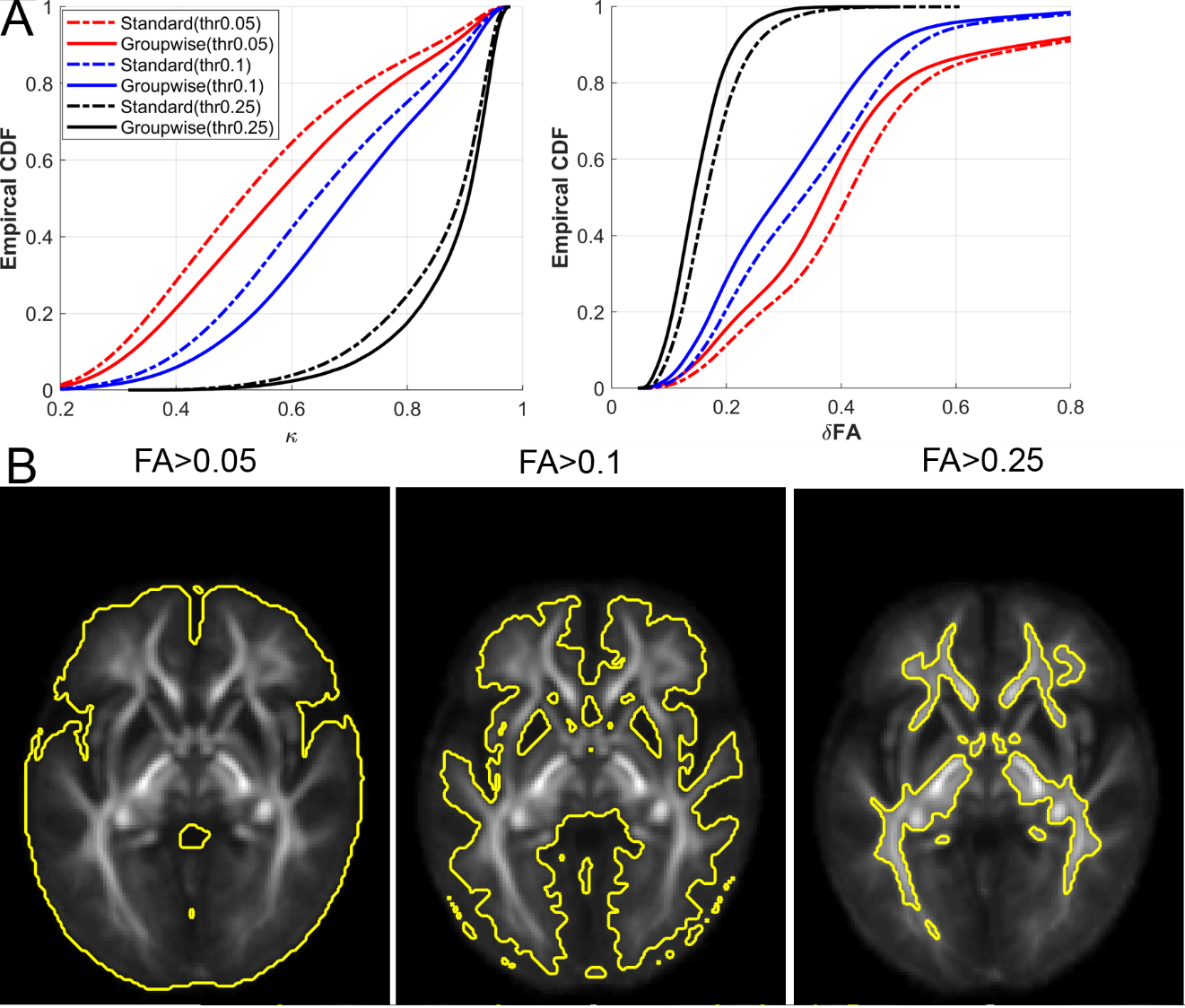

**Figure. S5**. Effects of brain mask on the performance of standard and groupwise tensor-based registration. (A) CDF plots of dyadic coherence (κ), and σ_FA_ of groupwise tensor-based registration (solid lines) and standard tensor-based registration (dotted lines) summed over the voxels within brain masks thresholded at FA > 0.05, FA > 0.1 and FA > 0.25, respectively. Regardless of the brain mask threshold used, groupwise tensor-based registration consistently outperformed standard tensor-based registration. (B) The enclosed brain regions for the brain masks thresholded at the three levels (0.05, 0.1 and 0.25) on the mean FA image.

***Effects of increasing number of iterations in groupwise tensor-based registration.*** Increasing the number of iterations to refine the registration during the affine and deformable transformation stages in the groupwise tensor-based registration improves registration accuracy, but the effect is negligible when the number of iterations is greater than 2 (Fig. S6).

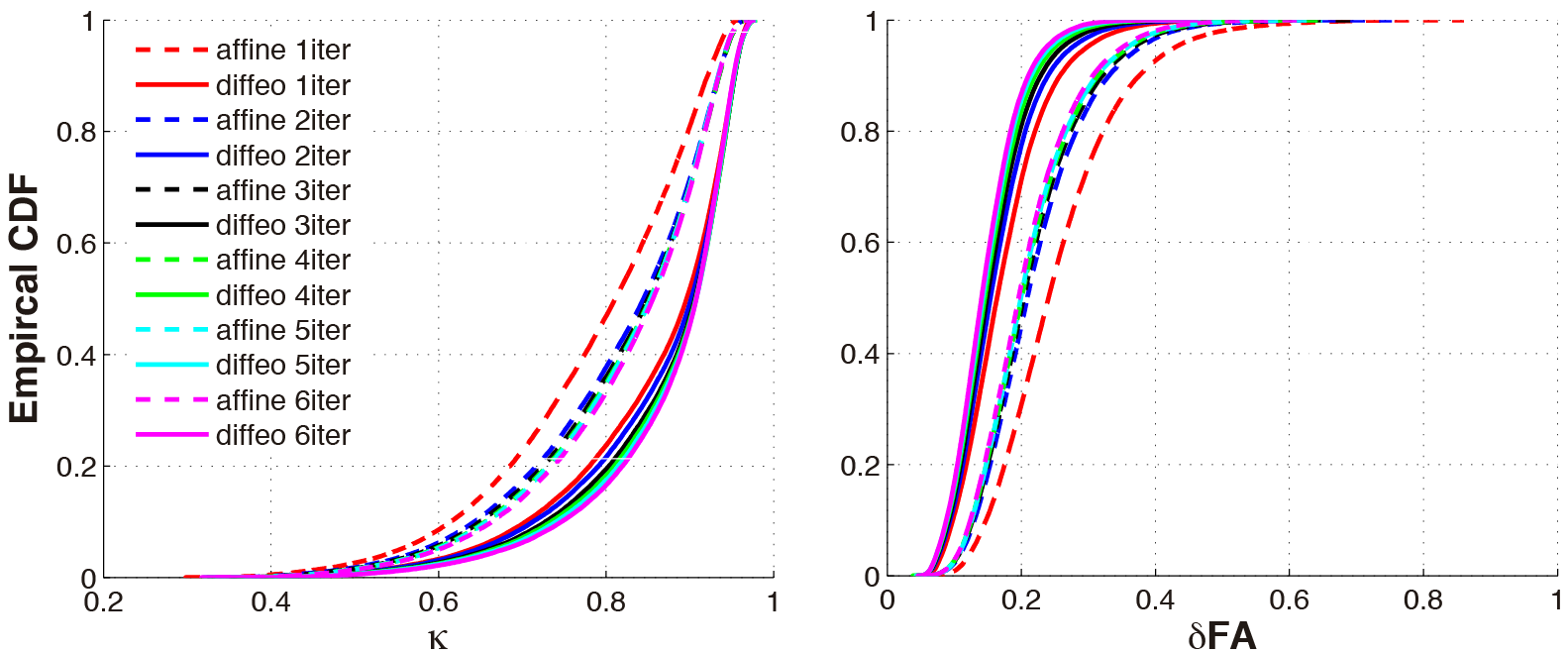

**Figure S6**. Effects of the number of iterations on the performance of groupwise tensor-based registration. By increasing the iteration number in the affine and deformable transformation stages from 1 (1iter) to 6 (6iter), the registration accuracy increases, but the improvement is trivial after the second iteration.

***Effect of choosing different images for generating the initial target template.*** We randomly selected two different DTI maps (with different brain sizes and brain shapes, see Fig. S7 (A) and Fig. S7 (B) for plots of their principal diffusion direction) with relatively clear tissue contrast and nudged them to closely match the origin of MNI space (the anterior commissure) and to be as straight as possible. The nudged DTI images were used as tensor templates for 6 dof rigid body transformation in the standard tensor-based registration (i.e., 6 dof rigid body transformation, followed by 12 dof affine transformation, and then diffeomorphic transformation) to generate the initial target template (the average of all aligned tensor images), separately. Fig. S7 (C) plots the principal diffusion direction of the initial target template generated by Fig. S7 (A). Fig. S7 (D) shows the principal diffusion direction of the initial target template generated by Fig. S7 (B). We can observe that the resulting initial target templates had the same brain size and shape even when the tensor templates for 6 dof rigid body transformation had different brain sizes and brain shapes.

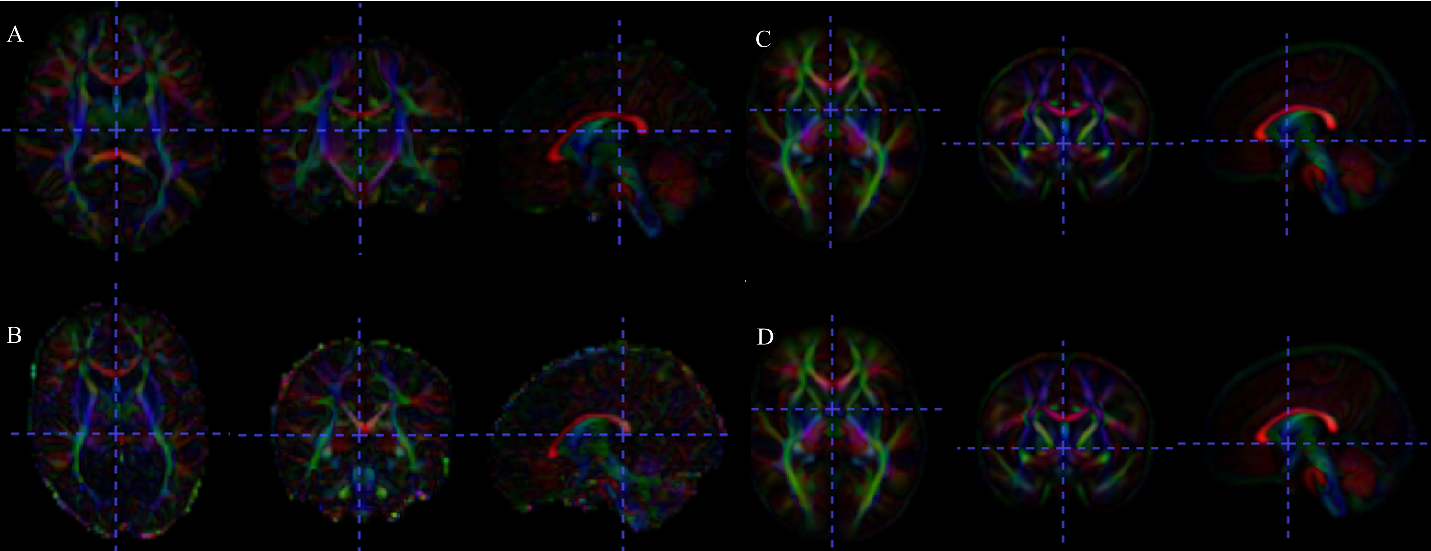

**Figure. S7**. Effect of choosing different tensor templates for 6-dof rigid body transformation to generate the initial target template using standard tensor-based registration. Principal diffusion directions of (A-B) two different tensor templates for 6 dof rigid body transformation, and (C-D) the resulting initial target template. Red, green and blue colors in the tensor map indicate the principal diffusion directions are in the left-right, anterior-posterior and inferior-superior orientations.
